# Long-range memory of growth and cycle progression correlates cell cycles in lineage trees

**DOI:** 10.1101/373258

**Authors:** Erika E Kuchen, Nils Becker, Nina Claudino, Thomas Höfer

**Affiliations:** German Cancer Research Center (DKFZ), 69120 Heidelberg, Germany; Bioquant Center, University of Heidelberg, 69120 Heidelberg, Germany

## Abstract

Mammalian cell proliferation is controlled by mitogens. However, how proliferation is coordinated with cell growth is poorly understood. Here we show that statistical properties of cell lineage trees – the cell-cycle length correlations within and across generations – reveal how cell growth controls proliferation. Analyzing extended lineage trees with latent-variable models, we find that two antagonistic heritable variables account for the observed cycle-length correlations. Using molecular perturbations of mTOR and MYC we identify these variables as cell size and regulatory license to divide, which are coupled through a minimum-size checkpoint. The checkpoint is relevant only for fast cell cycles, explaining why growth control of mammalian cell proliferation has remained elusive. Thus, correlated fluctuations of the cell cycle encode its regulation.

---

Cells of the same type growing in homogeneous conditions have variable cycle lengths (*1*). However, the mechanisms that set the duration of the cell cycle remain poorly understood. To first approximation, cycle lengths may be explained by the progression of the regulatory machinery of the cell cycle through a series of checkpoints in the presence of molecular noise (e.g., due to transcription (*2–4*). However, the cell cycles within a lineage tree are correlated in a non-intuitive pattern. Cycle lengths are similar in symmetrically dividing sister cells, which may be due to the inheritance of molecular regulators across mitosis (*5–9*). Ancestral correlations in cycle length fade rapidly, often disappearing already at the grandmother or even the mother cell. Nevertheless, the cycle lengths of cousin cells are correlated, indicating that the grandmother cell exerts an effect through at least two generations. These high intra-generational correlations in the face of weak ancestral correlations have been observed in cells as diverse as bacteria (*10*), cyanobacteria (*11*), lymphocytes (*12*) and mammalian cancer cells (*13,14*). Theory shows that more than one heritable factor is required to generate such correlations (*12*), one of which has been proposed to be the circadian clock (*14,15*). However, the identity of these memory conferring factors has not been probed experimentally.

To elucidate the origins of multiple memories in cell lineage trees, we identified candidate mechanisms by Bayesian model selection and tested them in molecular perturbation experiments. We began by asking how far intra-generational cell-cycle correlations extend within lineage trees. To this end, we generated extensive lineage trees by imaging and tracking TET21N neuroblastoma cells for up to ten generations during exponential growth (Fig. 1A and fig. S1A). Autonomous cycling of these cells is controlled by ectopic expression of the MYC-family oncogene *MYCN*, overcoming the restriction point and thus mimicking the presence of mitogenic stimuli (*3*). The distribution of cycle lengths (Fig. 1B and fig. S1B) was constant throughout the experiment (Fig. 1C and fig. S1C) and similar across lineages (Fig. S1D), showing absence of experimental drift and of strong founder cell effects, respectively. To determine cycle-length correlations without censoring bias caused by finite observation time (Fig. S2A) (14), we truncated all trees after the last generation completed by the vast majority (> 95%) of lineages. The resulting trees were 5–7 generations deep, enabling us to reliably calculate Spearman rank correlations between relatives up to second cousins (Fig. 1D and fig. S2B). Cycle-length correlations of cells with their ancestors decreased rapidly with each generation (Fig. 1E). However, the correlations increased again when moving down from ancestors along side-branches—from the grandmother to the first cousins and also from the great-grandmother to the second cousins (Fig. 1E). The intra-generational correlations (first and second cousins) were significantly larger than what would be expected from inheritance of cycle length alone (*13*), e.g., by passing on cell-cycle regulators (Supplemental Text). The discrepancy between theoretical expectation and experimental data was not due to spatial inhomogeneity or temporal drift in the data (Fig. S2, C-E). Thus, the lineage trees show long-ranging intra-generational correlations that cannot be explained by the inheritance of cell-cycle length.

**Fig. 1.**
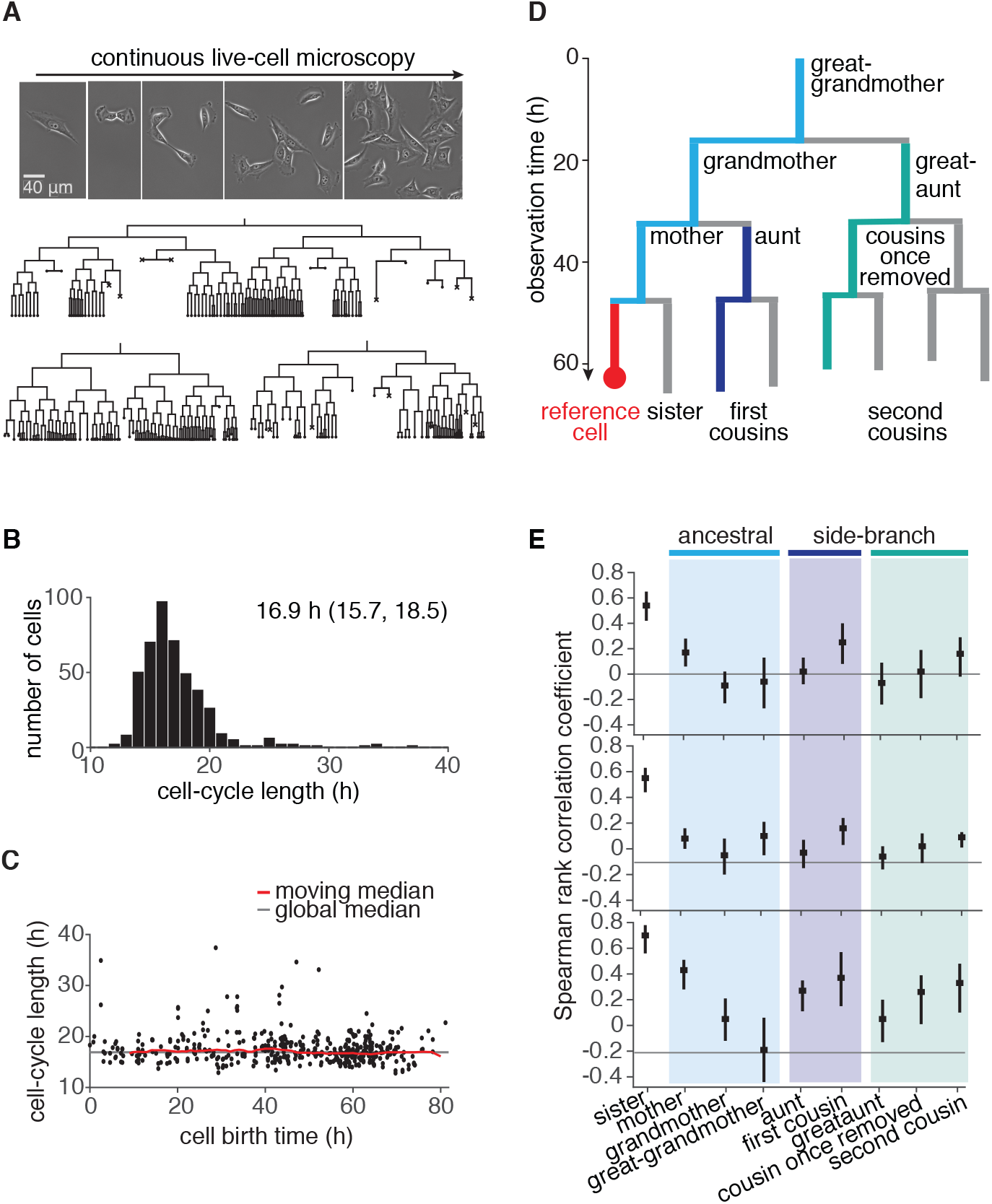
Cell-cycle lengths and their correlations captured by live-cell imaging. (**A**) Live-cell microscopy of neuroblastoma TET21N cell lineages. Sample trees shown with cells marked that were lost from observation (dot) or died (cross). (**B**) Distribution of cycle lengths, showing median length and interquartile range. (**C**) Cycle length over cell birth time shows no trend over the duration of the experiment. (**D**) Lineage tree showing the relation of cells with a reference cell. (**E**) Spearman rank correlations of cycle lengths between relatives (with bootstrap 95%-confidence bounds) of three independent microscopy experiments. Color code as in D.

We used these data to search for the minimal model of cell-cycle control that accounts for the observed correlation pattern of lineage trees (Supplementary Text). To be unbiased, we assumed that cycle length is controlled jointly by a yet unknown number *d* of cellular quantities ***x*** = (*x*_1_,…, *x_d_*), modeled as Gaussian latent variables. The cycle length is determined by the sum over these variables with positive weights *α_l_*, 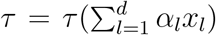. In any given cell *i*, ***x***^*i*^ is composed of an inherited component, determined by ***x*** in the mother, and a cell-intrinsic component that is uncorrelated with the mother. The inherited component is specified by an inheritance matrix **A**, such that the mean of ***x***^*i*^ conditioned on the mother’s ***x*** is 〈***x***^*i*^|***x***〉 = **A*x*** (Fig. 2A). The cell-intrinsic component causes variations around this mean with covariance 〉(***x***^*i*^–**A*x***)(***x***^*i*^–**A*x***)^⊤^|***x***〉 = **I**, where, with appropriate normalization of the latent variables, **I** is the unit matrix. Additional positive correlations in sister cells may arise due to inherited factors accumulated during, but not affecting, the mother’s cycle (*7–9*); additional negative correlations may result from partitioning noise (*16*). These are captured by the cross-covariance between the intrinsic components in sister 1 and 2, 〈(***x***^1^ – **A*x***)(***x***^2^ – **A*x***)^⊤^|***x***〉 = *γ***I**. In total, *d*(*d* + 1) parameters can be adjusted to fit the correlation pattern of the lineage trees: the components *a_lm_* of the inheritance matrix **A**, the weights *α_l_* and the sister correlation *γ*. Together, these inheritance rules specify bifurcating first-order autoregressive (BAR) models for multiple latent variables governing cell-cycle duration (generalizing previous work (*17*)).

**Fig. 2.**
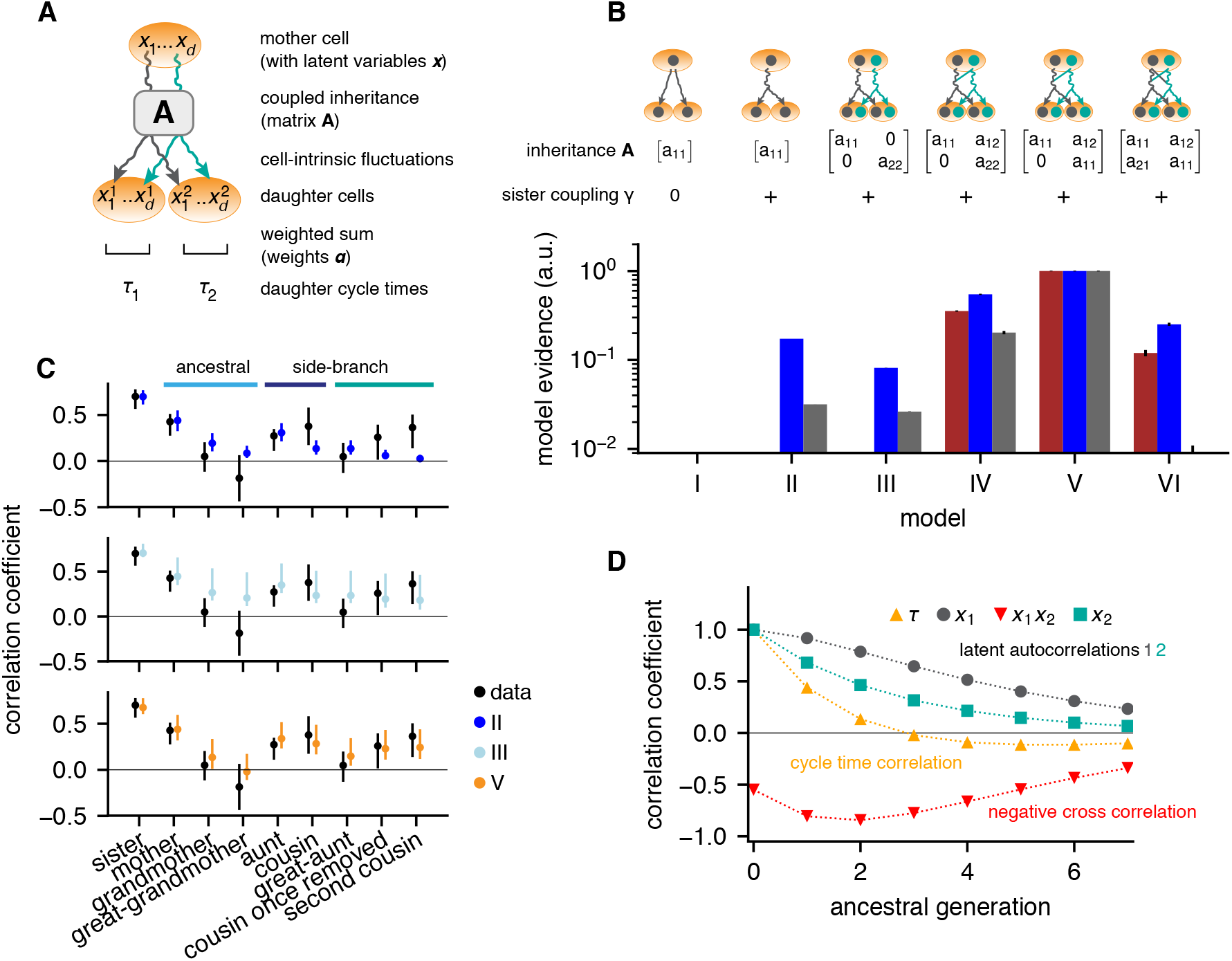
Bifurcating autoregressive inheritance models. (**A**) Coupled inheritance of *d* Gaussian latent variables *x_l_* and cell-intrinsic fluctuations generate cycle lengths. (**B**) Relative model evidences calculated for *d* = 1,2, for the indicated inheritance matrices **A** = [*a_lm_*] and sister coupling *γ*. Model V with unidirectionally coupled inheritance best explains the data. Error bars from Monte-Carlo integration. (**C**) Single-variable inheritance (Model II) fails to generate strong intra-generational correlations; uncoupled inheritance (III) fails to generate low ancestral correlations; Model V fits the data best. Rank correlations of the data shown with bootstrap 95%-confidence bounds. Model prediction bands were generated as the range of the parameter sets with likelihood higher than 15% of the best fit, corresponding to a Gaussian 95% credible region. (**D**) Model V, best-fit autocorrelation function along an ancestral line for cycle lengths τ and latent variables. Long-range memory in the latent variables is anticorrelated and masked in observed cycle times.

We evaluated the likelihood of the measured lineage trees and used it to rank BAR models of increasing complexity according their support by the experimental data, expressed as Bayesian evidence (Fig. 2B). The simplest model that generated high intra-generational correlations was based on the independent inheritance of two latent variables (Model III; Fig. 2C, cyan dots), whereas one-variable models failed to meet this criterion (Model II, Fig. 2C, blue dots and Model I). However, Model III consistently overestimated ancestral correlations and hence its relative evidence was low (< 10% for all data sets). We then accounted for interactions of latent variables. The most general two-variable model, allowing for bidirectional interaction (Model VI), overfitted the experimental data and consequently had low evidence. The highest support from the data was achieved by Models with unidirectional coupling, such that *x*_2_ in the mother negatively influenced *x*_1_ inherited by the daughters (with *a*_12_ < 0 and *a*_21_ = 0; Fig. 2B, Models IV and V; fig. S3, Model VII). Among these, the parsimonious model with only one self-inheritance parameter for both variables (*a*_11_ = *a*_22_ > 0) was preferred (Model V, Fig. 2B and 2C, orange dots). This preferred Model V produced a remarkable inheritance pattern (Fig. 2D): Individually, both latent variables had long-ranging memories, with 50% decay over 2–3 generations. However, the negative unidirectional coupling cross-correlated the variables negatively along an ancestral line, resulting in cycle length correlations that essentially vanished after one generation. Nevertheless, strong intra-generational correlations were reproduced by the model due to long-range memories of latent variables together with positive sister-cell correlations (*γ* > 0). We conclude that the coexistence of rapidly decaying ancestral correlations and extended intra-generational correlations is explained by the inheritance of two latent variables, one of which inhibits the other.

In order to divide, cells need a minimum size and license to progress through the cell cycle from the regulatory machinery. This regulatory license is realized through multiple bistable checkpoints (*18*). Mammalian cells continue to grow in size when regulatory license is withheld (e.g., in the absence of mitogens) (*19*) and growth is not otherwise constrained (e.g., by mechanical force or growth inhibitors) (*20*). As cell size and regulatory license are fundamental for cell-cycle progression and subject to inheritance, we asked whether these two quantities underlie the two latent variables in the BAR model. Indeed, there is an inbuilt negative effect of cell-cycle regulation in the mother cell on the time required for growth in the daughter cell: If regulatory license for cell division is delayed and the mother cell grows large, its daughters will be large at birth, grow to the critical size quickly and hence could have shorter cell cycles. Conversely, very short cell cycles generated this way will have to be followed by longer ones to allow cell size to recover. To test this idea quantitatively, we developed a simple model of growth and cell-cycle progression on cell lineage trees. We introduced the variables cell “size” *s*, measuring the metabolic, enzymatic and structural resources accumulated during growth, and *p*, characterizing the progression of the cell-cycle regulatory machinery. Unlike the latent variables of the BAR model *x*_1_ and *x*_2_, their mechanistic counterparts *s* and *p*, respectively, are governed by rules reflecting basic biological mechanisms (Fig. 3A), as follows. Size *s* grows approximately exponentially (*16,21*) and is divided equally between the daughters upon division. The progression variable *p* determines the time taken for the regulatory machinery to complete the cell cycle, which is controlled by the balance of activators and inhibitors of cyclin-dependent kinases. These regulators are inherited across mitosis (*5, 7–9*) and hence the value of *p* is passed on to both daughter cells with some noise. Cells divide when they have exceeded a critical size, requiring time *τ_g_*, and the regulatory machinery has progressed through the cycle, which takes an approximately log-normally distributed time (*3, 6*) modeled as *τ_p_* = exp(*p*). Hence the cycle length is *t* = max(*τ_g_, τ_p_*). Apart from requiring a minimal cell size for division, the model does not implement a drive of the cell cycle by growth and thus allows cells to grow large during long cell cycles. We fitted this model to the measured lineage trees by Approximate Bayesian Computation (Fig. S4A). The parameterized model yielded a stationary cell size distribution (Fig. S4B) and reproduced the cycle-length distribution (Fig. 3B, fig. S4C) as well as the ancestral and intra-generational correlations (Fig. 3C, fig. S4D). Thus the dynamics of cell growth and cell-cycle progression, coupled only through a minimal-size checkpoint, explain the intricate cycle-length patterns in lineage trees.

**Fig. 3.**
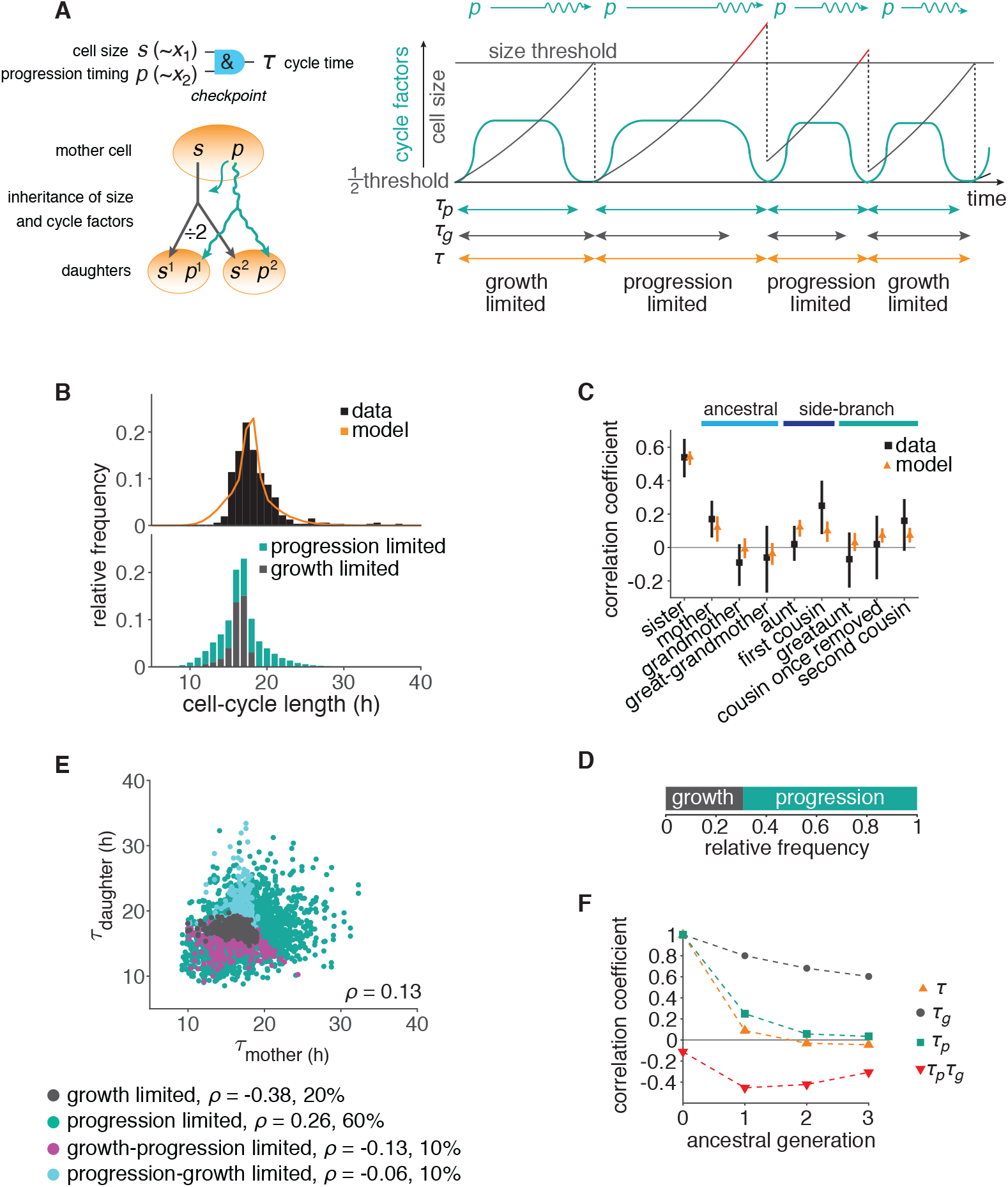
The growth-progression model. (**A**) Scheme of the growth-progression model with heritable variables relating to cell size s and cycle progression timing *p*. (**B**) Measured and simulated cell-cycle length distributions (upper). Model distribution resolved by the division-limiting process (lower). (**C**) Measured and modeled correlation pattern with Spearman rank correlation coefficient and bootstrap 95%-confidence bounds. (**D**) Proportion of simulated cells limited by growth or progression. (**E**) Correlation of simulated mother-daughter cycle lengths colored by their division-limitation: both by *τ_g_* (gray), both by *τ_p_* (green), mother *τ_p_* - daughter *τ_g_* (magenta), mother *τ_g_* - daughter *τ_p_* (cyan). Percentage of cells in each subgroup and their correlation coefficients are shown. (**F**) Autocorrelations along ancestral line of cycle length τ, growth time *τ_g_*, the progression time *τ_p_* and their cross-correlation *τ_p_τ_g_*.

If this growth-progression model captures the key determinants of the cell-cycle patterns in lineage trees, it should be experimentally testable by separately perturbing growth versus cell-cycle progression. We first derived model predictions for these experiments. Individual cell cycles in the model are either growth-limited, when division happens upon reaching the minimal size, or progression-limited, when cells grow beyond the minimal size until the cycle is completed (Fig. 3D and fig. S4E). Strictly positive cycle-length correlations between mothers and daughters arose when both generations had progression-limited cell cycles, whereas mother-daughter pairs in which at least one member was growth-limited reduced these correlations (Fig. 3E). The decorrelating effect of the coupling between cell-cycle progression and cell growth was also seen by the negative cross-correlations between these two processes along ancestral lines (Fig. 3F and fig. S4F), as in the BAR model (cf. Fig. 2D). Thus both proliferation regulation and cell growth exhibit longer-ranging memories whose inheritance across generations is masked by their negative coupling. Using the model to simulate perturbation experiments, we predict that growth inhibition will increase intra-generational correlations relative to ancestral (mother-daughter) correlations whereas slowing cell-cycle progression will decrease these correlations (Fig. 4A).

**Fig. 4.**
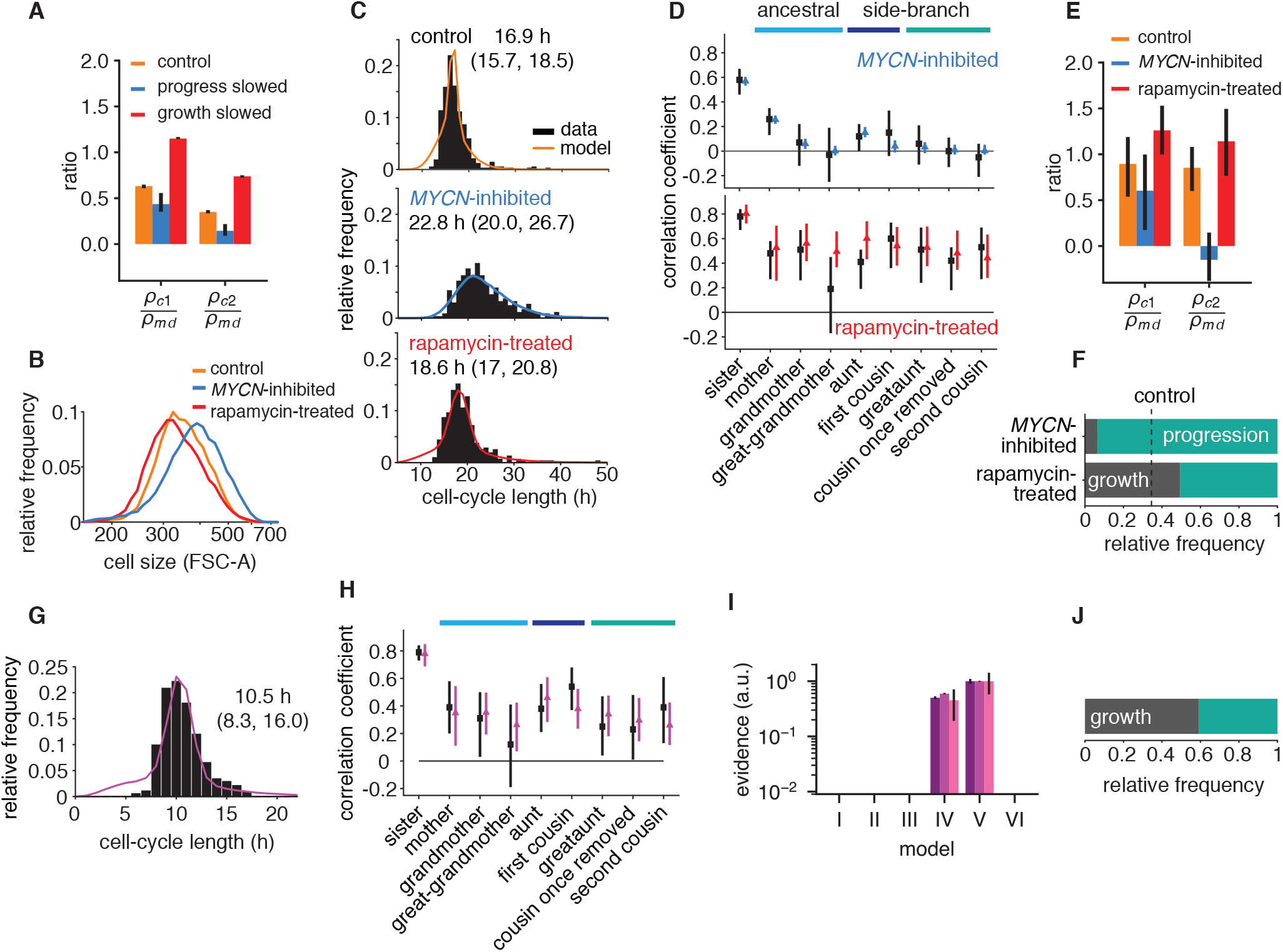
Targeted perturbation of growth and cell-cycle progression. (**A**) Predictions for changes in the ratio *ρ_c_/ρ_md_* of cousin to mother-daughter correlations, when slowing growth or cycle progress compared to the best-fit parameters (control). *ρ*_*c*1_ = first cousins, *ρ*_*c*1_ = second cousins. (**B–F**) Experimental perturbations of cycle progress and growth by MYCN inhibition and rapamycin treatment, respectively. (B) Cell size distribution. Areal forward scatter measured experimentally by flow cytometry for control *high*-MYCN, *MYCN*-inhibited and rapamycin-treated TET21N neuroblastoma cells; n=2, representative experiment shown. (C) Measured and best-fit model cycle length distributions. Median and interquartile range are indicated. (D) Measured (black) and best-fit correlation pattern of MYCN-inhibited and rapamycin-treated cells with Spearman rank correlation coefficient and 95%-confidence bounds. (E) Measured cousin / mother-daughter correlation ratios. (F) Proportion of simulated cells limited by growth or progression, using best-fit parameters for MYCN inhibition or rapamycin-treatment. (**G–J**) Embryonic stem cell data provided by Filipczyk et *al*. (*22*) re-analyzed for cell-cycle correlations. (G) Cycle length distribution of data (black) and growth-progression model (purple). (H) Measured (black) and modeled (purple) correlation pattern using the growth-progression model. (I) Model evidences of the BAR model, version numbering as in Fig. 2B. (J) Proportion of simulated cells limited by growth or progression as in F.

To test this prediction, we slowed cell-cycle progression experimentally by reducing MYCN, exploiting the doxycycline-tunable *MYCN* gene integrated in the TET21N cells. Cells grew to larger average size (Fig. 4B, blue line) over longer and more variable cell cycles (Fig. 4C). These data show that lowering MYCN slowed cell-cycle progression without compromising growth. Further consistent with this phenotype, expression of mTOR, a central regulator of metabolism and growth (*19*), was not lowered (Fig. S5A). In a separate experiment, we inhibited cell growth. To this end, we applied the mTOR inhibitor rapamycin, which reduced cell size (Fig. 4B, red line). This treatment also lengthened the cell cycle (Fig. 4C) but without changing MYCN protein levels (Fig. S5B). Thus lowering MYCN and inhibiting mTOR are orthogonal perturbations that act on cell-cycle progression and cell growth, respectively. As predicted by the growth-progression model, these perturbations resulted in markedly different cycle-length correlation patterns within lineage trees (Fig. 4D, E and fig. S5C, D): Lowering MYCN decreased intra-generational correlation and, in particular, removed second-cousin correlations. By contrast, rapamycin treatment strongly increased intra-generational correlations and caused ancestral correlations to decline only weakly. Collectively, these findings support the growth-progression model of cell-cycle regulation.

We then asked, conversely, whether the cycle-length correlation patterns experimentally observed in lineage trees contain information about the underlying regulation. To this end, we fitted the growth-progression model to MYCN and rapamycin perturbation data and again obtained good agreement with the data (Fig. 4C, D and fig. S5C-E). The fit parameters imply that the vast majority of cell cycles upon *MYCN* inhibition were progression-limited (Fig. 4F and fig. S4F), consistent with *MYCN* inhibition reinstating bistable restriction point control (*3*). By contrast, rapamycin treatment significantly increased the fraction of growth-limited cell cycles. Thus, the cycle-length correlation pattern in lineage trees reveals how strongly cell growth controls proliferation.

Finally, we analyzed time-lapse microscopy data of non-transformed mouse embryonic stem cells (*22*) that proliferate much faster than the neuroblastoma cells (Fig. 4G and fig. S5F). Side-branch correlations of cycle length increased with each generation from the common ancestor (Fig. 4H and fig. S5G), as seen in all previous data except for the *MYCN*-inhibited cells. Interestingly, the strength of the intra-generational correlations was most similar to the much more slowly dividing rapamycin-treated cells (cf. Fig. 4D). As before, the BAR model required two negatively coupled variables to account for these data (Fig. 4I, fig. S6). Fitting the growth-progression model to the data (Fig. 4G, H), we found that the majority (~60%) of cell cycles were limited by growth (Fig. 4J, fig. S5H), indicating that cycle length of fast proliferating mammalian cells is, to a large extent, controlled by growth.

In summary, our findings provide an unexpected link between two lines of research: The study of correlation patterns in cell lineage trees (*10,13,14*) and size regulation of the mammalian cell cycle (*19–21,23,24*). Using perturbations of cell growth and cell-cycle progression, we show that the non-intuitive correlation patterns of cycle length in lineage trees are explained by inheritance of cell size and cell-cycle regulators. Inheritance of both factors generates extended intra-generational correlations in cycle length, which are masked in ancestral lines by the decorrelating impact of cell-cycle progression on growth.

A quantitative model derived from the experimental data shows only weak interaction of size and regulatory license through a minimum-size checkpoint. This mode of regulation differs fundamentally from yeast, where growth drives progression (*25,26*). Indeed, in mammalian cells growth control becomes apparent only for very rapid cell cycles (e.g., once in ten hours in embryonic stem cells) or upon growth inhibition. This finding may explain why the regulation of the mammalian cell cycle by growth has been difficult to study experimentally. Since inheritance of regulators of the cell cycle and cell size across cell division are universal phenomena, we expect that correlation patterns will provide a quantitative tool to study their interaction across a wide range of organisms.

## Acknowledgments

We thank Frank Westermann and Tatjana Ryl for TET21N cells and laboratory setup; Timm Schroeder, Fabian Theis and Carsten Marr for embryonic stem cell data and discussions; the Hofer group for discussions. Grant support from BMBF (MYC-NET 0316076A and Sysmed-NB 01ZX1307), EU (FP7-HEALTH-2010 ASSET 259348), BMBF and EU (EraCoSysMed OPTIMIZE-NB 031L0087A) is gratefully acknowledged; T.H. is a member of CellNetworks.

## Supplementary materials

Supplementary Text Materials and Methods Fig. S1 - S6 References (*27–32*)

## References

1. J. A. Smith, L. Martin, Proceedings of the National Academy of Sciences of the United States of America 70, 1263 (1973).

2. S. Kar, W. T. Baumann, M. R. Paul, J. J. Tyson, Proceedings of the National Academy of Sciences of the United States of America 106, 6471 (2009).

3. T. Ryl, et al., Cell Systems 5, 237 (2017).

4. A. R. Barr, F. S. Heldt, T. Zhang, C. Bakal, B. Novák, Cell systems 2, 27 (2016).

5. S. Spencer, et al., Cell 155, 369 (2013).

6. S. Mitchell, K. Roy, T. A. Zangle, A. Hoffmann, Proceedings of the National Academy of Sciences of the United States of America 115, E2888 (2018).

7. H. W. Yang, M. Chung, T. Kudo, T. Meyer, Nature 549, 404 EP (2017).

8. A. R. Barr, et al., Nature Communications 8, 14728 (2017).

9. M. Arora, J. Moser, H. Phadke, A. A. Basha, S. L. Spencer, Cell reports 19, 1351 (2017).

10. E. O. Powell, Journal of general microbiology 18, 382 (1958).

11. Q. Yang, B. F. Pando, G. Dong, S. S. Golden, A. van Oudenaarden, Science (New York, N.Y.) 327, 1522 (2010).

12. J. F. Markham, C. J. Wellard, E. D. Hawkins, K. R. Duffy, P. D. Hodgkin, Journal of the Royal Society Interface 7, 1049 (2010).

13. R. G. Staudte, M. Guiguet, M. C. d’Hooghe, Journal of theoretical biology 109, 127 (1984).

14. O. Sandler, et al., Nature 519, 468 (2015).

15. N. Mosheiff, et al., Physical Review X 8, 021035 (2018).

16. Y. Sung, et al., Proceedings of the National Academy of Sciences 110, 16687 (2013).

17. R. Cowan, R. Staudte, Biometrics 42, 769 (1986).

18. B. Novak, J. J. Tyson, B. Gyorffy, A. Csikasz-Nagy, Nature cell biology 9, 724 (2007).

19. D. C. Fingar, S. Salama, C. Tsou, E. Harlow, J. Blenis, Genes & Development 16, 1472 (2002).

20. I. Conlon, M. Raff, Journal of biology 2, 7 (2003).

21. A. Tzur, R. Kafri, V. S. LeBleu, G. Lahav, M. W. Kirschner, Science 325, 167 (2009).

22. A. Filipczyk, et al., Nature Cell Biology 17, 1235 EP (2015).

23. M. B. Ginzberg, et al., eLife 7, e26957 (2018).

24. R. Kafri, et al., Nature 494, 480 (2013).

25. A. A. Amodeo, J. M. Skotheim, Cold Spring Harbor Perspectives in Biology 8, a019083 (2016).

26. G. Facchetti, F. Chang, M. Howard, Current Opinion in Systems Biology 5, 86 (2017).

27. K. Boudt, J. Cornelissen, C. Croux, The Gaussian Rank Correlation Estimator: Robustness Properties, SSRN Scholarly Paper ID 1689465, Social Science Research Network, Rochester, NY (2010).

28. E. Jaynes, Physical Review 106, 620 (1957).

29. J. Lin, A. Amir, Cell Systems (2017).

30. D. J. C. MacKay, Information Theory, Inference and Learning Algorithms (Cambridge University Press, 2003).

31. W. Lutz, et al., Oncogene 13, 803 (1996).

32. E. Meijering, O. Dzyubachyk, I. Smal, Methods in Enzymology 504, 183 (2012).

